# Irreversible temperature gating in trpv1 sheds light on channel activation

**DOI:** 10.1101/251124

**Authors:** Ana Sánchez-Moreno, Eduardo Guevara-Hernández, Ricardo Contreras-Cervera, Gisela Rangel-Yescas, Ernesto Ladrón-de-Guevara, Tamara Rosenbaum, León D. Islas

## Abstract

Temperature activated TRP channels or thermoTRPs are among the only proteins that can directly convert temperature changes into changes in channel open probability. In spite of a wealth of information, including several experimentally determined structural models of TRP channels, the mechanism of temperature activation remains unknown. We have carefully characterized the repeated activation of TRPV1 by thermal stimuli and discovered a previously unknown inactivation process, which is irreversible. This inactivation is associated with specific conformational changes in the membrane proximal domain. We propose that this form of gating in TRPV1 channels is a consequence of the heat absorption process that leads to channel opening.

## Introduction

Thermo TRP channels are unique among ion channels in that they can be activated by changes in temperature alone (Latorre et al., 2009). In particular, several members of the TRPV family respond to different ranges of high temperatures and contribute to the physiological sensation of temperature changes in mammals and other organisms (Gracheva et al., 2010; Gracheva et al., 2011).

The biophysical response of TRPV1 to temperature change is a large increase in current, produced both by an increase in the channel open probability and in the single channel-conductance, with the open probability showing the larger temperature dependence. The mechanism by which these channels accomplish the conversion of the absorbed heat into a conformational change that eventually triggers channel opening remains largely unknown (Castillo et al., 2017).

Nonetheless, some physical principles involved in the activation of heat-activated thermoTRP channels have been identified. A large positive enthalpy change in the order of tens of kcal·mol^-1^ are required for such activation, and is strictly balanced by a large entropy change. This enthalpy-entropy matching is necessary to maintain a large value of the associated rate constants for opening and thus ensure rapid channel activation (Yao et al., 2010a). It is also known, for TRP channels, that the heat dependence is mostly associated with channel opening and that the closing transition is not highly heat-dependent (Liu et al., 2003; Yao et al., 2010a).

Along with the thermodynamic constrains of activation, these channels behave as allosteric proteins in which several activating stimuli are coupled to their opening. In particular, allosteric activation of TRPV1 channels is indicated by the presence of multiple open and closed states (Liu et al., 2003). This allosterism, is also observable as a coupling between distinct activation pathways when channels are activated by more than one stimulus (Ahern et al., 2005). This has led to the adoption of quantitative allosteric formalisms as the preferred way of explaining gating in TRPV1 (Brauchi et al., 2004; Cao et al., 2013b; Jara-Oseguera and Islas, 2013; Jara-Oseguera et al., 2016).

However, the conformational changes responsible for channel gating by temperature are unknown. In contrast with the voltage sensor of voltage-gated channels, a “temperature sensor” has not been found and it has been suggested that temperature might affect several regions of the channel protein at once, perhaps explaining why so many regions have been implicated in temperature gating in thermoTRP channels (Brauchi et al., 2006; Grandl et al., 2008; Yao et al., 2010b, 2011; Cui et al., 2012). An attractive mechanism for channel gating by temperature that invokes a large difference in heat capacity between the closed and open conformations has been proposed (Clapham and Miller, 2011; Chowdhury et al., 2014). One possible way of realizing this mechanism is the exposure of hydrophobic or hydrophobic regions or residues to a hydrophilic or hydrophobic environment, respectively. These changes need not represent large conformational rearrangements.

In this report, we investigated the repeated activation of TRPV1 channels by temperature. We find that activation is coupled to a previously undidentifed temperature-dependent inactivation and show that this transition is irreversible. Our data is suggestive of a mechanism that might be the result of partial or complete unfolding of one or more regions of the channel protein during heat absorption.

## Results

### Activation of TRPV1 channels by temperature ramps

One of the hallmarks of TRPV1 channels is their high temperature-sensitivity, implying that high enthalpic changes are associated with channel gating. At the same time, heat induced activation is very rapid. Using fast IR-laser induced temperature changes, the activation rate constant has been estimated to be in the ms or sub-ms range (Yao et al., 2009). In this work, we made use of an alternate and previously reported method to apply controlled ramps of temperature to achieve reliable temperature activation of TRPV1 (Islas et al., 2015). Since our temperature ramps change with a rate of ∼15°C/s, and activation of TRPV1 by square temperature jumps has a time constant in the order of ms our measurements can be considered to be in quasi steady-state.

Figure 1A (bottom) shows the time course of a typical temperature ramp, measured 27 μm from the surface of the heater, where the recording pipette and membrane patch are also placed for experiments. Under these conditions, temperature changes in an almost completely symmetrical fashion (black, bottom). The resulting temperature-activated current through TRPV1 is shown in the same figure (top). TRPV1-mediated current can be activated at both negative (yellow trace) and positive (blue trace) potentials. Channel activity can be observed even at 20°C in multichannel patches and the increase in channel activity visibly occurs around 40°C (Figure 1B). Current activates and deactivates steeply, but in spite of the symmetric heat stimulus provided by the ramp, the current response is not symmetrical to heating (blue) and cooling (gray) (Figure 1C). The asymmetry in current activation and deactivation can be seen more clearly when the current is plotted as a function of ramp temperature (Figure 1D). It is apparent that the current generated upon temperature decrease (gray) is less steep than the up-ramp current (blue), so current activation-deactivation is accompanied by a high degree of hysteresis. Hysteresis is not a consequence of the speed of the ramp since activation by a slower ramp (7 °C/s) produces the same result (Figure supplement 1). A less temperature-dependent deactivation current could in part be explained because current closure upon cooling is almost temperature-independent (Yao et al., 2010a). However, in this study it will be shown that it reflects irreversible changes in temperature-dependence.

**Figure 1.**
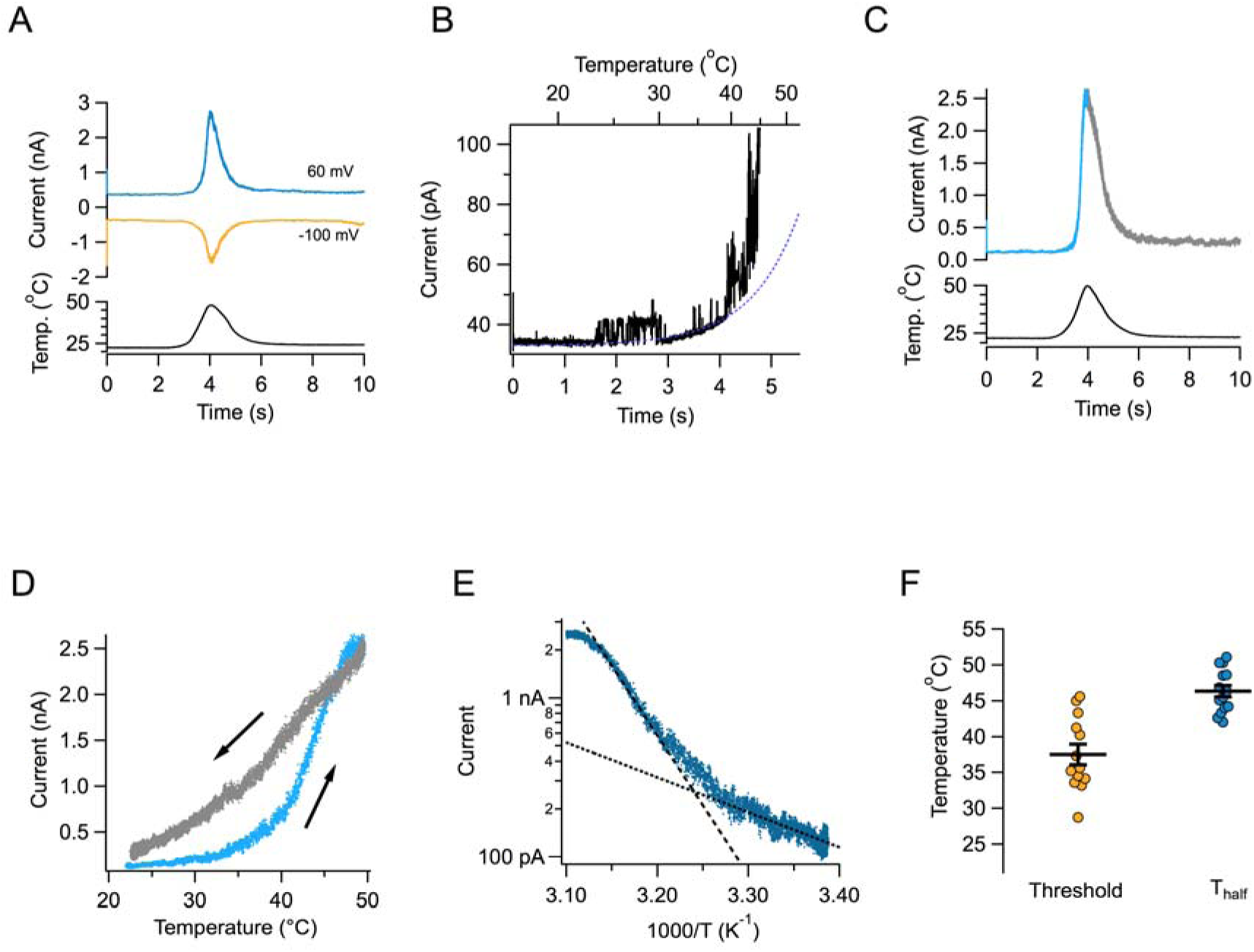
Activation of TRPV1 current by temperature ramps. (A) A ramp of total duration 4 s (2 up and 2 down) is shown below the currents activated in an inside-out patch at two different voltages. The maximum temperature reached was 47.8°C. (B) A different patch showing the up and down regions of the temperature ramp in different color. Notice the asymmetry in the kinetics of the current. (C) Single channels can be observed in a patch containing hundreds of TRPV1 channels. In this patch, the detectable increase in open probability happens around 40°C. (D) The current in panel (B) is plotted as a function of the temperature ramp. The arrows indicate the direction of temperature increase (blue) or decrease (gray). It is evident that the paths followed by activation and deactivation are different. (E) Van’t Hoff representation of the current (teal) as a function of inverse temperature. The threshold of current activation is determined from the intersection of the two exponential functions representing the leak current (black dotted line) and the TRPV1 current (black dashed line). (F) Activation properties of TRPV1 by heat. Threshold and temperature of half– activation (T_half_) for 14 patches.

Transformation of the activating region of the current to a van’t Hoff plot, allowed for the estimation of the enthalpic change (ΔH) involved in opening the channels as the slope of an exponential function fitted to the steep part of the activation curve. For the current in Figure 1e the value is 40 kcal∙mol^-1^. Although channel activation by temperature does not exhibit a threshold (Figure 1B), some researchers have defined an operative threshold as the intersection of an exponential function describing channel activation and an exponential defining the temperature dependence of leak (patch seal) current (Cui et al., 2012). Figure 1f shows the values of threshold defined in this fashion, along with the temperature at which half of the maximum current is obtained (T_half_) for several patches. These values are similar to figures previously reported (Yao et al., 2010a).

### TRPV1 inactivates with repetitive activation

A key novel observation while repetitively activating TRPV1 by reproducible temperature ramps is that the magnitude of the current is reduced with each ramp. Strikingly, after several activations TRPV1-mediated currents completely disappear (Figure 2A). The loss of current becomes more dramatic if currents are activated at larger temperatures. When channels are activated by temperatures near 55°C, current decay is visible even before the ramp reaches its maximum temperature (Figure 2B). The decay of current as a function of time is temperature-dependent. Current loss when activating by ramps that reach moderate temperatures (41-43°C) proceeds more slowly than when channels are activated by ramps above 50°C (Figure 2C). Current loss is not caused by patch excision, since it can be observed in whole-cell recordings and in outside-out recordings, with similar characteristics (Figure supplement 2). This suggests that the mechanism of current loss is a property of the TRPV1 polypeptide and is not caused by the loss of a stabilizing factor upon patch excision, as described for other TRP channels (Liu and Qin, 2005; Rohács et al., 2005; Klein et al., 2008).

**Figure 2.**
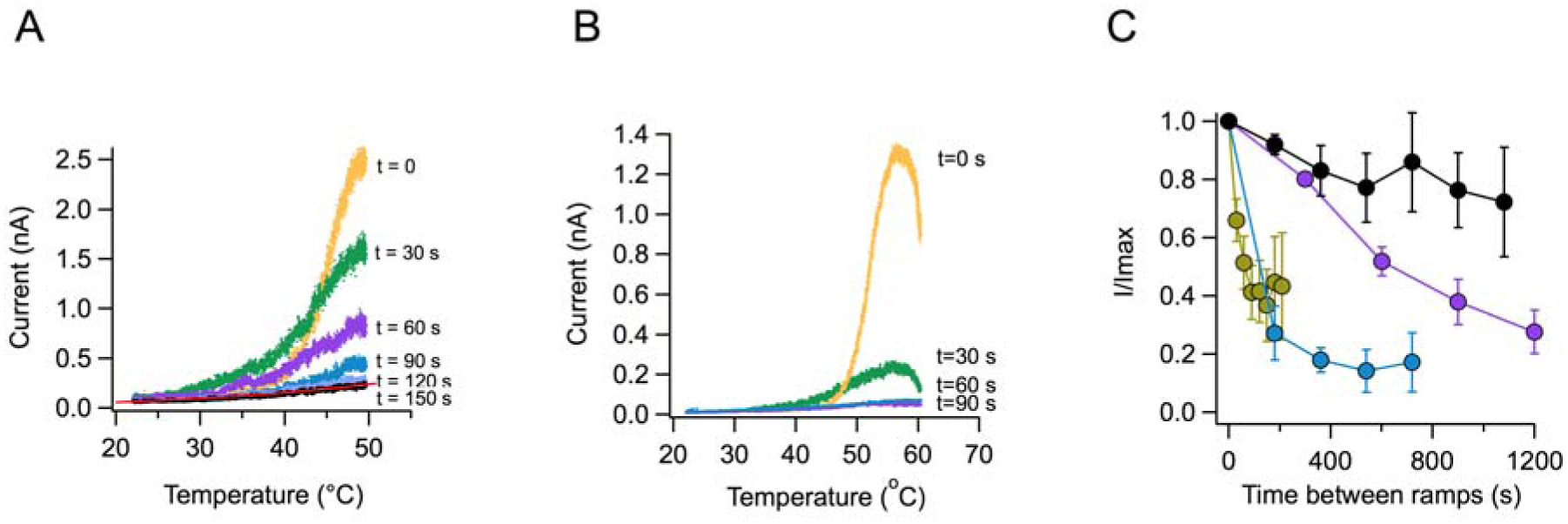
Loss of current with repetitive activation in TRPV1. (A) Activation (up) part of the response to a temperature ramp in an inside-out patch expressing TRPV1 channels. The maximum temperature attained was 49.7°C. Ramps of total (up and down) duration of 4 s were applied every 30 s. Note the marked reduction of peak current as well as the decreased slope of activation. The trace at 150 s is essentially the response of the patch, since all TRPV1 current has been lost. This trace is fitted to an exponential dependence of temperature with an associated enthalpy of 14 kcal∙mol^-1^. (B) Response of a different patch to a ramp of higher temperature (peak temperature 59°C). The loss of current is evident even during the up ramp before the ramp has reached its maximum temperature. The response to subsequent ramps is much more diminished at this higher temperature. (C) Time course of current loss as a function of different maximum ramp temperatures. Maximum temperatures attained during the activation ramps are: Black circles, 41-43°C, purple circles, 47-49°C, blue circles, 51-53°C, lemon circles, 51-53°C, with a shorter interval between ramps.

### Thermal inactivation of TRPV1 is irreversible

In order to further characterize current loss with repetitive thermal stimuli, currents were inactivated by more than 50 % and then reactivated after a recovery period. Surprisingly, current could not be elicited after a recovery period of up to two minutes. Lack of recovery is observed when current is inactivated by application of several ramps at moderate temperature (Figures 3A and C) or during a single ramp at high temperature (Figures 3D and F). Since current loss is highly temperature dependent, we investigated if low temperature could help in the recovery process. Again, current loss was irreversible even if the temperature was lowered to ~ 10°C during the recovery period (Figures 3B, C, E and F).

**Figure 3.**
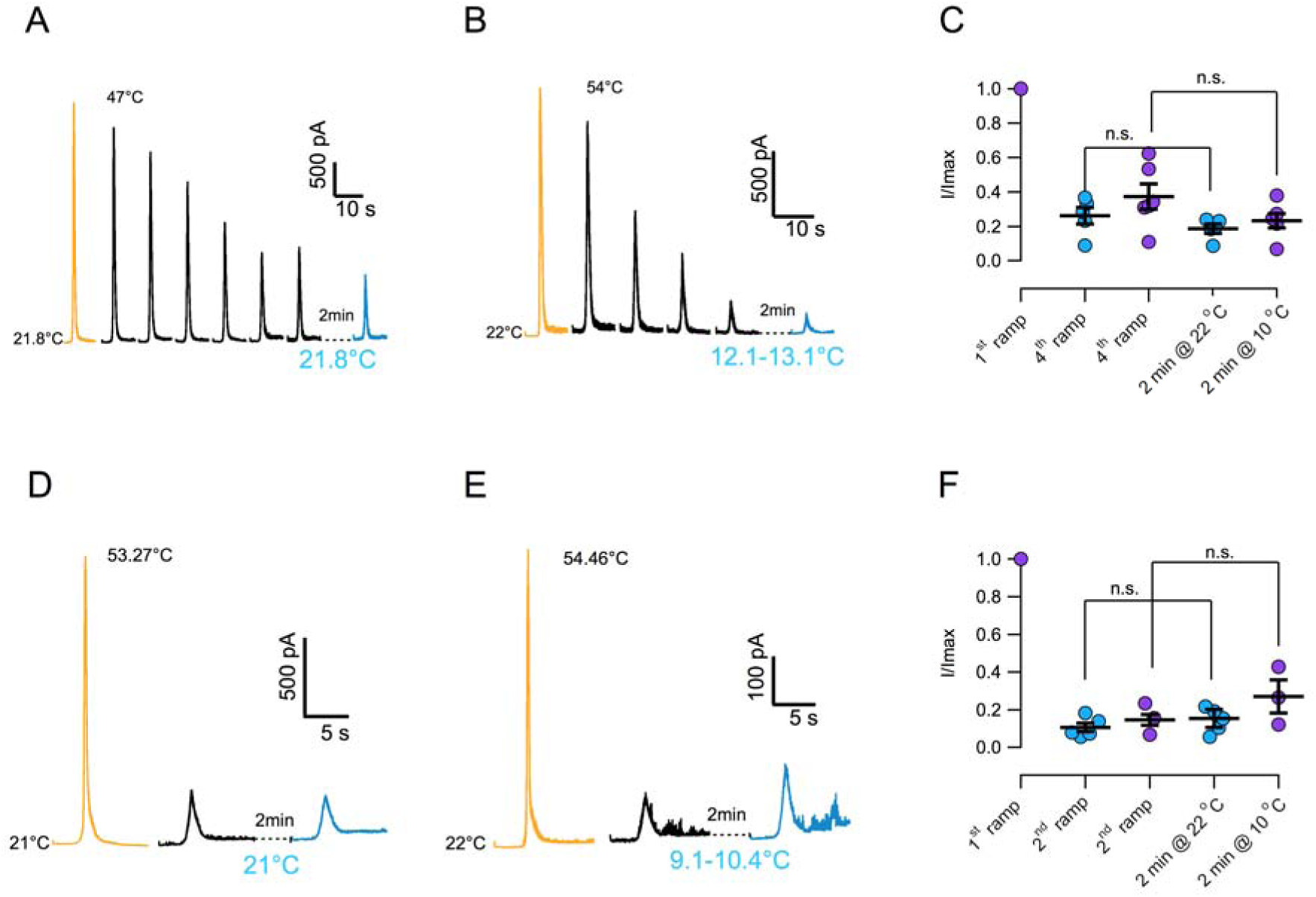
Inactivation of TRPV1 by multiple temperature ramps is irreversible. (A) In this particular patch, seven consecutive ramps to the temperature indicated were applied with an interval of 30 s to produce ~60% inactivation. The patch was left at the bath temperature (21.8 °C) for a 2 min recovery period, after which another activation ramp did not elicit a current larger than the residual current after inactivation, indicating absence of recovery from inactivation. (B) Another patch was subjected to 5 activation ramps that produced ~80 % inactivation. This time, during the 2 min recovery period, bath temperature was lowered to between 12-13°C. Again, no current was produced by an activation ramp, indicating no recovery. (C) Summary of results from several experiments indicating no recovery of current after a 2 min period at normal bath temperature (~22°C, blue circles) or in a cooled bath (10°C, purple circles). (D) Similar to (A) but in this patch the current was elicited by a ramp to a larger temperature, which produced a larger inactivation in a single ramp (80 %). After a 2 min period at 21°C, no current was recovered. (E) Same as in (C) but with a recovery period at 10°C. No recovery of current is evident. (F) Summary of results from several experiments as in (D) and (E) indicating no recovery of current after a 2 min period at normal bath temperature (~22°C, blue circles) or in a cooled bath (10°C, purple circles). Absence of significant recovery was based on a Student t-test with p values indicated in the figure. The label n.s. indicates non-significance. All p-values were much greater than 0.01.

An important observation is the fact that the reduction of current amplitude resulting with repetitive stimulation is also accompanied by a dramatic change in the sensitivity of channels to temperature. This sensitivity is determined by enthalpic (ΔH) and entropic (ΔS) components (Voets et al., 2004). In order to estimate the magnitude of ΔH and ΔS, we fitted activation curves with a simplified temperature-dependent allosteric model (Figure 4A and Methods). The response of TRPV1 to the first ramp results in a very steep activation curve, with average enthalpy of 72 kcal∙mol^-1^. Subsequent activation curves show reduced slopes, indicating a reduction of the ΔH and ΔS associated with channel opening (Figure 4B). Currents activated by the second ramp, applied 30 s after the first, shows a ~30 % reduction of ΔH and after 5 ramps with maximum temperature of 48.7°C, the enthalpy associated with activation is reduced to to ~10 kcal∙mol^-1^, or a Q_10_ value of 1-2, which is similar to diffusion processes (Sidell and Hazel, 1987). When ΔH and ΔS are plotted for several patches for the first and second ramp, the reduction in ΔH and ΔS is obvious and a striking enthalpy-entropy correlation is observed, which has been shown before to apply to TRPV1 (Yao et al., 2010a), and which remains valid even for channels that undergo irreversible inactivation (Figure 4C).

**Figure 4.**
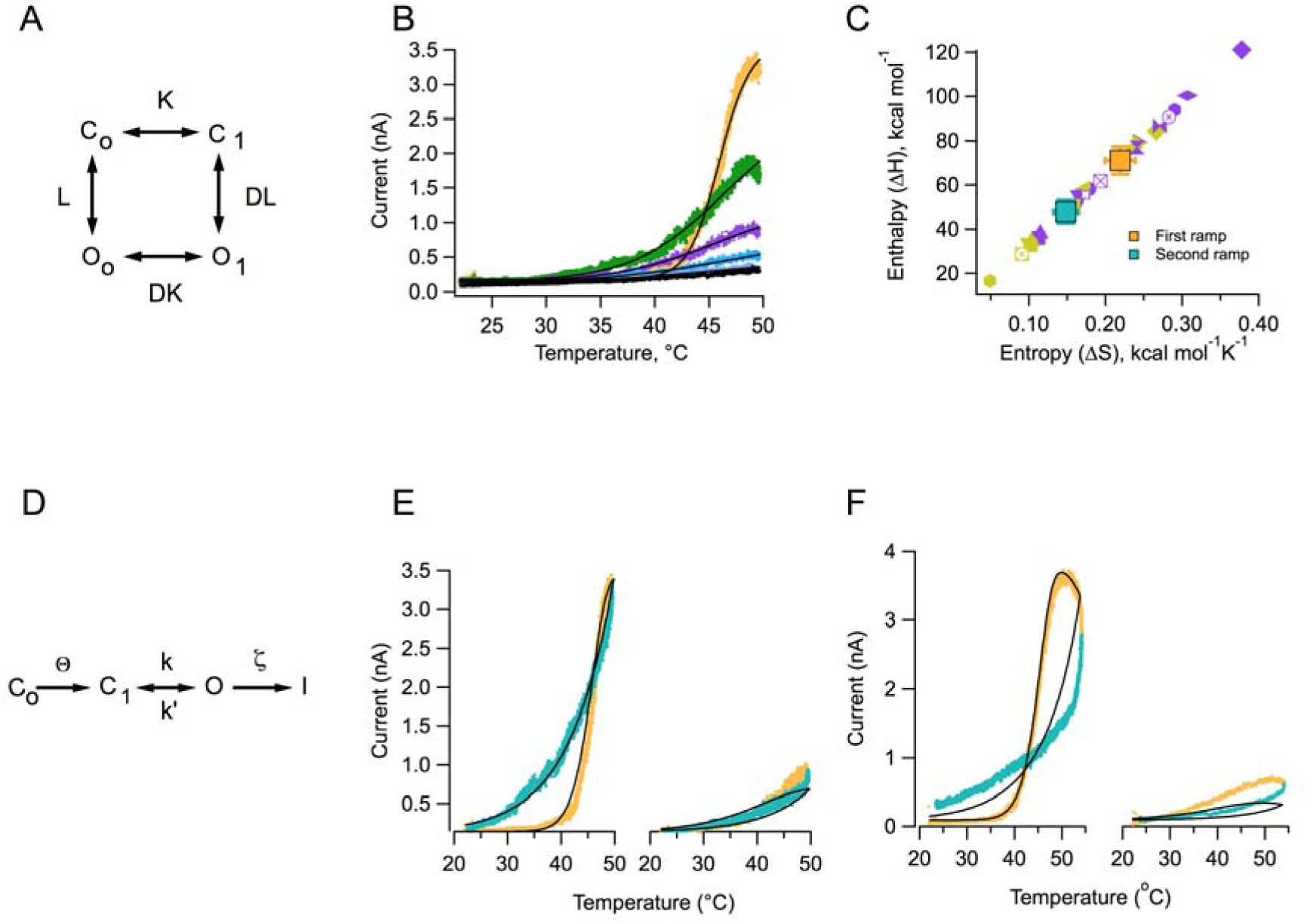
Thermal activation models recapitulate the characteristics of activation and inactivation by heat. (A) Allosteric activation model used to obtain estimates of enthalpy (ΔH) and entropy (ΔS) changes during activation. Definition of variables of the model is given in the Methods section. (B) Black continuous curves are fits of the allosteric model to activation functions obtained from successive ramps to 49.6°C at 30 s intervals. The value of D was fixed to 830 and the value of L_o_ was fixed to 0.001. The respective values of ΔH and ΔS for each time interval are: t=0, 134.96 kcal∙mol^-1^, 0.422 kcal∙mol^-1^ K^-1^; t=30 s, 44.20 kcal∙mol^-1^, 0.137 kcal∙mol^-1^ K^-1^; t = 60 s, 39.49 kcal∙mol^-1^, 0.122 kcal∙mol^-1^ K^-1^; t=90 s 25.94 kcal∙mol^-1^, 0.079 kcal∙mol^-1^ K^-1^; t=120 s, 24.18 kcal∙mol^-1^, 0.074 kcal∙mol^-1^ K^-1^. (C) The values of ΔH and ΔS from 14 patches for the first (purple symbols) and second ramps (lemon symbols). Each patch is represented by a different symbol. The average values of ΔH and ΔS are shown by the yellow and teal squares for the first and second ramp, respectively. Error bars are s.e.m. (D) A linear activation model with two irreversible conformational changes. The rate constants are temperature-dependent and are defined in the Methods section (C_i_, closed states, O open state and I inactivated state). Up and down currents in response to a first ramp to 49.6°C (left) and the same ramp 60 s later (right). The black lines are the predicted open probabilities expected from the model in (D) for two symmetrical ramps applied at the same rate of temperature change as the experimental ramp and separated by 60 s at 20°C. The values of variables in the model are: θ_θ_ = 0.4×10^7^ s^-1^, ΔH_θ_ = 74.88 kcal∙mol^-1^, ΔS_θ_ = 0.209 kcal∙mol^-1^K^-1^, θ_k_ = 0.5x10^6^ s^-1^, ΔH_k_ = 37.44 kcal∙mol^-1^, ΔS_k_ = 0.0942 kcal∙mol^-1^K^-1^, θ_k’_ = 0.03 s^-1^, ΔH_k’_ = 13.2 kcal∙mol^-1^ and ΔS_k’_ = 0.06 kcal∙mol^-1^K^-1^, θ_ζ_ = 0.5x10^6^ s^-1^, ΔH_ζ_ = 40 kcal∙mol^-1^. (F) Same as in (E), but for currents elicited at a higher maximum ramp temperature of 55°C. The parameters of the model are the same as for the simulations in (e).

In order to obtain a quantitative representation of the phenomenology discussed above, we applied a simple sequential model that incorporates irreversible activation and inactivation steps. The first activating transition is steeply temperature-dependent but irreversible, followed by a reversible step with smaller associated ΔH and ΔS, leading to channel opening. After channels open, they enter an absorbent inactivated state. This transition is also temperature dependent (Figure 4D). The model nicely reproduces the highly hysteretic shape of the currents activated by the up and down phases of a ramp to 49°C, the reduced temperature dependence of activation after the first ramp and also reproduces the reduction of current observed upon repetitive stimulation (Figure 4E). The same model with the same parameters also reproduces the behavior of currents when the ramp is taken to a peak temperature of 54.2°C, although the fit is less good (Figure 4F). These results suggest that TRPV1 channels indeed undergo an irreversible conformational change upon heat absorption and channel opening and subsequently enter an absorbing inactivated state.

Up to here, we have shown that current loss by repetitive stimulation is irreversible and is accompanied by a dramatic reduction in the ability of the channels to respond to heat. In order to evaluate the extent of the conformational change accompanying current loss, we performed experiments to determine if current loss only represents an inability of the channel to be reactivated by temperature or if it is a more widespread phenomenon, by examining capsaicin activation after current loss. Surprisingly, activation by capsaicin is also impaired after multiple activation ramps (Figure 5). Before heat activation, capsaicin activates robust currents, but the response to a second application of capsaicin after the patch has undergone several activating ramps, is highly diminished (Figure 5B). There is a clear correlation between the extent of current loss induced by temperature and the magnitude of current activated by the second application of capsaicin (Figure 5C), suggesting that the remaining capsaicin-sensitive channels have not yet undergone inactivation.

**Figure 5.**
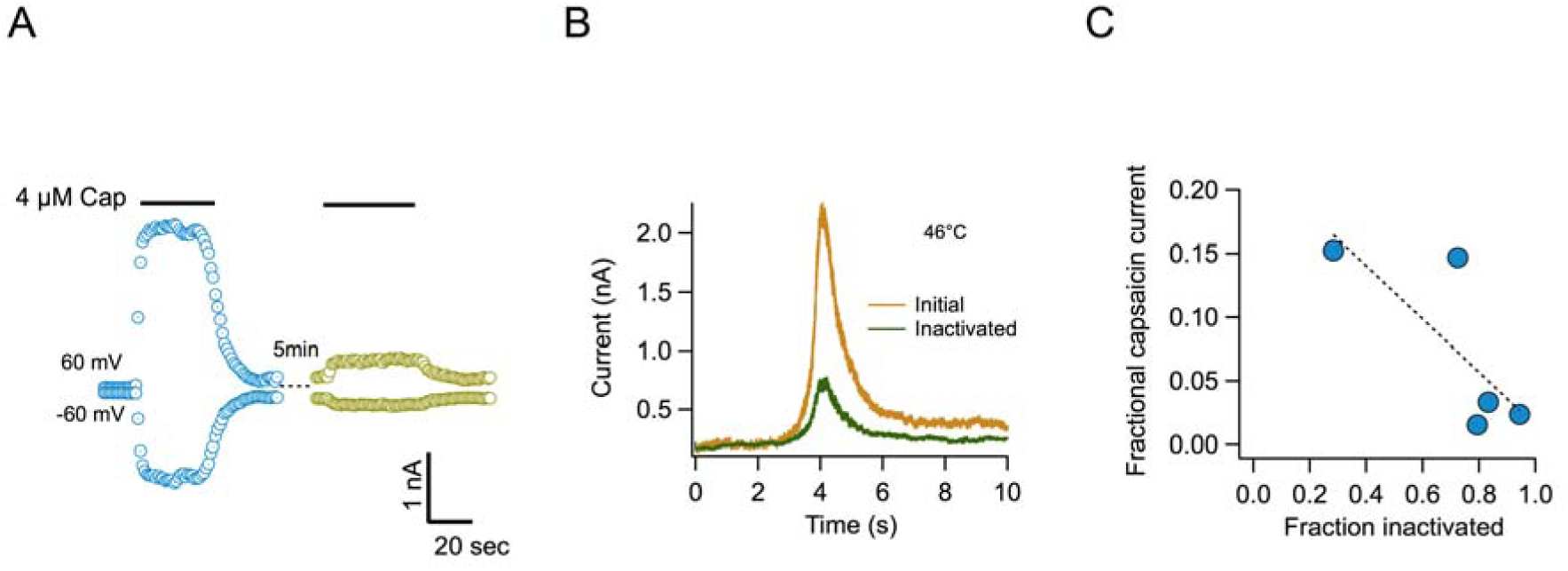
Inactivation by temperature renders the channel irresponsive to activation by capsaicin. (A) Response of channels in an inside out patch to application of a saturating concentration of capsaicin before (blue) and after (lemon) application of 4 activation ramps to the indicated temperature. Observe the much-reduced response to capsaicin application. (B) Currents in response to the first and fourth temperature activation ramps, applied between the two applications of capsaicin in (A), showing the reduction of current after the fourth ramp. (C) Inverse correlation between the degree of inactivation of TRPV1 currents and the response to reapplication of 4 μM capsaicin. The dotted line is a fitted linear function through the data, with a correlation coefficient, r = -0.77.

Irreversible current loss or inactivation suggests that heat activation causes an important conformational change in TRPV1. It has been shown that the membrane proximal linker domain (MPD) contains two cysteine residues, C386 and C390, in close proximity (yellow, Figure 6A), which can be induced to form a disulphide bond by oxidizing reagents. This disulphide results in partial activation of current through the channel (Chuang and Lin, 2009). We used this form of activation as a reporter of the conformational state of the channel. Control patches in which 10 µM of the oxidizing agent 5,5’ – dithiobis (2-nitrobenzoic acid) (DTNB) was applied to the inside of the membrane showed activation of current in the absence of capsaicin, which was about a third the size of the capsaicin-activated current in the same patch (Figure 6B). When currents were activated by temperature ramps after being transiently activated by capsaicin, subsequent application of DTNB failed to elicit any current (Figure 6C). Notice that even a single temperature ramp produced a diminished response to DTNB (Figure 6D), suggesting that current loss might be coupled to the activation process. These results indicate that heat activation increases the separation between cysteines and/or the accessibility of at least one of them. Two cysteine-less channel constructs in which 15 (15 cys less) or 16 (16 cys less) out of 18 cysteines were removed (Salazar et al., 2008) do not respond to DTNB (Data not shown) and still retain heat activation, albeit with reduced sensitivity, but importantly, retain temperature-dependent inactivation, indicating that cysteines are not necessary for temperature-dependent activation or inactivation (Figure supplement 6).

**Figure 6.**
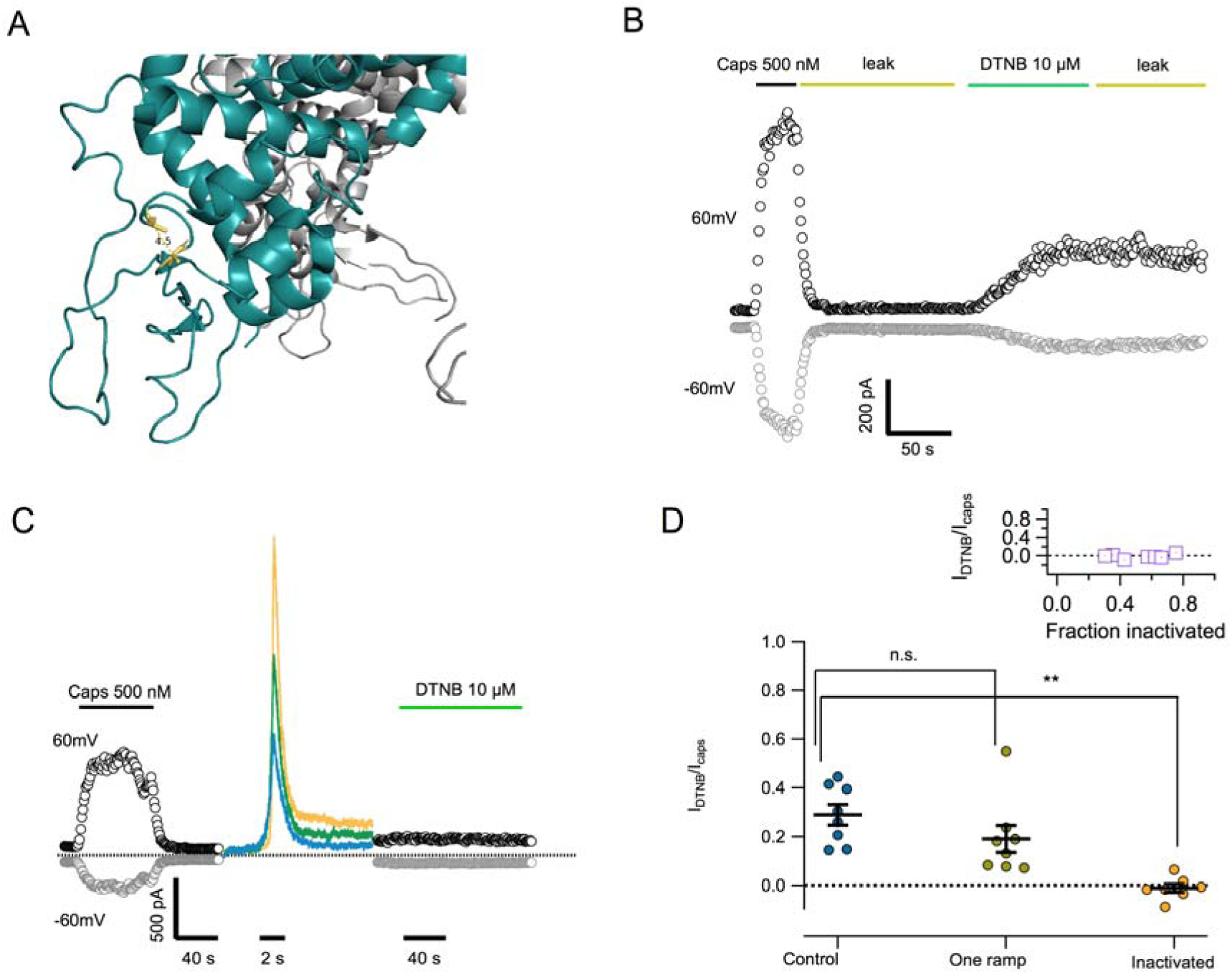
Inactivation is accompanied by a conformational change in the proximal linker domain. (A) Molecular model of the proximal linker domain of TRPV1 (pdb file:) showing the location of cysteines C386 and C390. The distance between the α-carbons of these cysteine residues is 4.5 Å. (B) Application of DTNB to inside out patches produces partial activation of TRPV1. The DTNB-activated current in this patch is 26% of the capsaicin-activated current. Washing off DTNB (leak) does not remove activation, indicating a covalent modification. (C) TRPV1 current in a patch was evaluated by application of to 500 nM capsaicin. Subsequently, three activation ramps to 51°C were applied, producing current loss. Application of DTNB after heat activation failed to produce measurable current. (D) Quantitation of the effect of heat activation on the activation of TRPV1 by DTNB. Disulphide bond formation induced by DTNB produces a fractional current of 0.28 ± 0.042 activation when compared to 500 nM capsaicin. After one thermal activation event (ramp to 46°C) this fraction is reduced to 0.19 ± 0.055 and when current loss is elicited by multiple temperature activation ramps, DTNB is incapable of activating any current, fractional current is -0.01 ± 0.017. The inset shows that there is no correlation between the degree of inactivation and the size of the DTNB-activated current. Statistically significant difference is indicated by ** and mean p < 0.01. The label n.s. indicates non-significance.

To find further support for the hypothesis that heat activation induces a conformational change that renders the channel non-responsive to activating stimuli, we applied a fluorescence-based approach to measure conformation of the channel. By incorporating non-canonical fluorescent aminoacids, a specific fluorescence signal can be recorded from proteins (Chatterjee et al., 2013; Lang and Chin, 2014). In this case, we first demonstrated that L-ANAP can be successfully incorporates at position 374 in rTRPV1 channels (Figure supplement 7) and that the fluorescence of the non-canonical aminoacid L-ANAP is quenched by the hydrophobic anion dipycrylamine (DPA), which can be intercalated in the plasma membrane. DPA quenches fluorescence of other fluorophores via a FRET mechanism (De-la-Rosa et al., 2013). We calculated a R_o_ for the ANAP-DPA pair of 4.1 nm. Quenching by DPA can be used to estimate distances and changes in distance between fluorophores in the channel protein and DPA in the plasma membrane (Chanda et al., 2005; Taraska and Zagotta, 2007).

We carried out experiments in which the amount of quenching by DPA in the absence of channel activation was compared to that observed after heating membrane sheets containing channels with ANAP incorporated at position 374 (Figure 7A). Position 374 lies at the end of a β-sheet that is almost continuous with the cysteines at positions C386 and C390 and thus we think should be a reporter of a conformational change in the same region. The measurements were carried out in deroofed HEK293 cells expressing ANAP-374 channels. Since the fluorescent protein mCherry was fused to the c-terminus of TRPV1-Y374TAG, mCherry fluorescence indicates full-length channels that by necessity incorporated ANAP. Membranes containing channels that incorporated ANAP where thus identified as regions with co-localization between mCherry and ANAP (Figure 7B). Correct incorporation of ANAP at position 374 was also confirmed by whole-cell recording of capsaicin-activated currents (Figure supplement 7), further indicating that fluorescence arises from functional channels. These membranes displayed correct spectra for ANAP and mCherry, indicating proper incorporation and protein folding (Figure 7C). DPA addition to these membrane sheets produces a 20 % reduction in the fluorescence of 374ANAP channels (Figure 7D, blue circles), indicating quenching.

**Figure 7.**
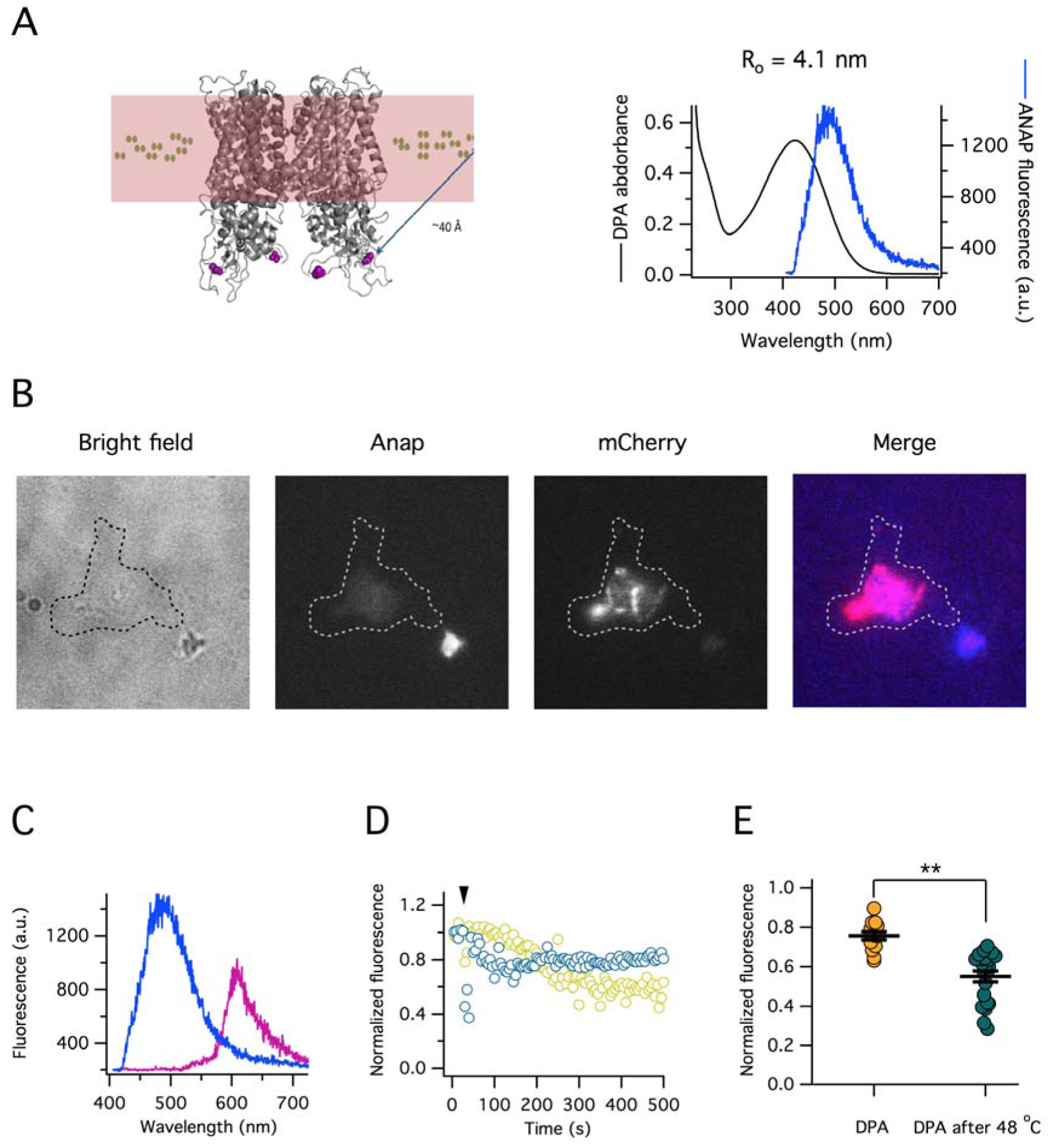
Inactivation is accompanied by a conformational change that brings the n-terminus closer to the plasma membrane. (A, left) Molecular model of TRPV1 showing the location of the Y374 residue (purple) and the DPA molecules in the membrane (green hexagons). (A, right) The fluorescence emission spectrum of ANAP and the absorption spectrum of DPA show a large region of overlap, rendering them a good FRET pair. The calculated R_o_ value for this pair is 4.1 nm. (B) Representative images of a membrane sheet expressing rTRPV1-374ANAP-mCherry channels. Fluorescence images are rendered in gray scale. The overlap of the ANAP and mCherry channels is shown in pinkish color, demonstrating overlap of the two signals, presumably arising from the same channels. (C) Emission spectra of the ANAP and mCherry regions of the same membrane sheet as in (B), demonstrating the correct spectra and showing correct folding of TRPV1mCherry incorporating ANAP. (D) Quenching of ANAP fluorescence by the incorporation of DPA into the membrane sheet. Addition of DPA is indicated by the arrowhead. Control experiment is shown by blue symbols, the temperature was 24°C throughout. Addition of DPA to a different membrane sheet after being held at 48°C for 2 min. The increased quenching of ANAP fluorescence is due to increased proximity to the membrane. The delay in the response is due to differences in the bath perfusion between experiments. (E) Results from 13 control membrane sheets and 22 membrane sheets heated to 48°C for 2 min. The means are statistically different (p < 0.001), which is indicated by **.

Membrane sheets expressing 374ANAP channels that were subjected to 46°C for 2 min, displayed increased quenching (40%, Figure 7D, lemon circles and Figure 7E), suggesting that heating produced a non-transient a conformational change in which 374ANAP is closer to the membrane, and confirming the presence of a conformational change associated to temperature-induced inactivation.

## Discussion

The molecular underpinnings of activation by temperature in thermoTRP channels remain largely unknown. In TRPV1 channels, several regions of the protein have been demonstrated to have effects in the amount of heat involved in opening the channel (Yao et al., 2011) or in the threshold for activation (Cui et al., 2012; Laursen et al., 2016). The pore region of TRPV1 has also been implicated in regulating thermal activation (Grandl et al., 2008; Grandl et al., 2010; Yang et al., 2010). Making use of a reproducible and reliable method of repetitive thermal stimulation, here we report on the existence of an inactivation process in TRPV1, which had not been previously described. Inactivation proceeds with a time course in the order of seconds, similar to the slow inactivation processes described in Kv channels (Hoshi et al., 1991; Liu et al., 1996). The time course is strongly dependent on the maximum temperature attained during the activation ramp, behaving as if inactivation was coupled to opening. In this regard, temperature-dependent inactivation in TRPV1 also resembles open state inactivation in voltage-dependent channels, particularly N-type inactivation in Shaker (Demo and Yellen, 1991) and fast inactivation in mammalian Na^+^ channels (Aldrich et al., 1983).

However, the apparent resemblance to classical ion channel inactivation mechanisms is superficial. This inactivation process in TRPV1 is irreversible and is accompanied by a marked reduction of the channel’s sensitivity to temperature. Furthermore, after temperature-dependent inactivation, the channels can no longer be activated even by other stimuli such as capsaicin application. A simple kinetic model can satisfactorily explain our experimental results. This model incorporates two irreversible transitions, one that is related to activation by heat and one that represents entrance to the inactivated state. Although highly simplified, this scheme can explain both, the reduction in enthalpic and entropic components of activation observed with repetitive stimulation and the reduction of current observed with time and with temperature. This model implies that heat is absorbed through multiple conformation transitions, but some transitions might involve molecular rearrangements that cannot be reversed.

Irreversible transitions in proteins are more generally associated with large conformational changes or denaturation reactions (Privalov et al., 1989). Some enzymes become irreversibly inactivated by elevating temperature (Ahern and Klibanov, 1985) through a mechanism that is not completely well defined, but involves deamidation of Asn and Gln residues, which in turn induce an irreversible conformational change. We show with two methods, cysteine modification and fluorescence quenching, that temperature dependent inactivation of TRPV1 is accompanied by a conformational change in the membrane proximal linker domain (MPLD), which along with other regions of the N-terminus of TRPV1, TRPV3 and TRPA1, has been implicated in thermal activation (Cordero-Morales et al., 2011; Yao et al., 2011; Liu and Qin, 2017). We have detected these conformational changes with a conventional cysteine modification method and through analysis of fluorescence from the non-canonical aminoacid, ANAP. Direct visualization of ANAP fluorescence during heating of membranes expressing channels where ANAP was incorporated was not possible due to the observation that ANAP fluorescence has very high dependence on temperature and imaging conditions are drastically altered during heating, producing large imaging artifacts. For this reason, we have adopted an end-point fluorescence quenching approach and demonstrate that ANAP can be quenched by DPA, a membrane intercalating hydrophobic anion.

We propose that our results are compatible with a unique sequence of events during heat absorption and conversion to pore opening. Heat might be absorbed in several regions of the channel, leading to partial contributions to channel opening, but some regions might absorb heat through a mechanism akin to partial polypeptide denaturation. One such regions seems to be the MPD, since we can detect a temperature–induced conformational change in it, but it might not be the only one. This partial denaturation leads to channels that undergo this transition becoming less able to absorb or convert heat into a conformational change, resulting in reduced enthalpy of activation. Finally, after opening, channels become sufficiently distorted to enter this novel inactivation state irreversibly. This is consistent with a previous proposal that processes with high heat capacity (∆C_p_) might be involved in thermoTRP channel activation (Clapham and Miller, 2011; Chowdhury et al., 2014). Some such processes are large conformational changes, which have not being observed (De-la-Rosa et al., 2013; Ruigrok et al., 2017) or alterations of hydrophobic interactions through solvent exposure, which might be what we observe here.

Why was this prominent inactivation process not reported previously? Careful perusal of current recordings available in the literature give hints that researchers had observed inactivation of TRPV1 current upon activation by temperature in Xenopus TRPV1 (Saito et al., 2016), mouse TRPV1(Cui et al., 2012) and rat TRPV1 (Cao et al., 2013a). Finally, this form of temperature-dependent inactivation should play a physiological role in regulating excitability of temperature-sensing neurons. Channels that are irreversibly inactivated are most likely subject to recycling (Sanz-Salvador et al., 2012). It will be interesting to see what is the role of this form of channel regulation in physiological conditions.

## Materials and Methods

### Cell Culture and transfection

HEK293 cells were grown on 100 mm culture dishes with 10ml of DMEM, Dulbecco´s Modified Eagle Medium (Invitrogen) containing 10 % fetal bovine serum (Invitrogen) and 100 Units/ml-100 μg/ml of penicillin-streptomycin (Invitrogen), incubated at 37°C in a 5.2 % CO_2_ atmosphere. When cells reached 90 % confluence, the medium was removed and cells were treated with 1 ml of 0.05 % Trypsin-EDTA (Invitrogen) for 5 min. Subsequently, 1 ml of DMEM with 10 % FBS was added. The cells were mechanically dislodged and reseeded in 30 mm culture dishes over 5x5 mm coverslips for electrophysiology or in 35 mm glass bottom dishes, for imaging. In both cases, 2 ml of complete medium were used. Cells at 70 % confluence were transfected with the appropriate construct as indicated, using jetPEI transfection reagent (Polyplus Transfection). For patch-clamp experiments, pEGFP-N1 (BD Biosciences Clontech) was cotransfected with rTRPV1 to visualize successfully transfected cells via fluorescence. Electrophysiological recording were done one or two days after transfection.

### DNA constructs and mutagenesis

Electrophysiological experiments were carried out on rat TRPV1-WT cloned in the plasmid pcDNA3. An rTRPV1-mCherry construct was used as template for introduction of the TAG codon for L-ANAP incorporation. mCherry was fused to rTRPV1 in the c-terminus after a 5’-GGSGGSGGS-3’ linker, using standard overlap PCR methods. This construct was cloned into the pcDNA3 expression vector. TAG mutants were prepared on this background using mutagenic primers and complete plasmid PCR amplification with the KOD Hot Start polymerase. After PCR, the samples were treated with DpnI nuclease and transformed into *E. coli* DH5α. Confirmation of correct mutagenesis as made through automatic sequencing.

### Current recording

Patch clamp recordings were made from HEK293 cells expressing TRPV1 in the inside-out, whole-cell and outside-out configurations of the patch-clamp recording technique. Inside-out and outside-out recordings were made using symmetrical solutions consisting of 130 mM NaCl, 10 mM HEPES, 5 mM KCl and 5 mM EGTA for calcium free conditions, pH 7.2 adjusted with NaOH. Whole-cell recordings of TRPV1-mCherry and Y374ANAP-mCherry channels were made using a bath solution with 130 mM NaCl, 10 mM HEPES, 0.5 mM CaCl_2_, pH 7.2 adjusted with NaOH and a pipette solution with 130 mM NaCl, 10 mM HEPES, 5 mM KCl and 5 mM EGTA.

Macroscopic currents were low pass filtered at 1 kHz, sampled at 20 kHz with an Axopatch 200B amplifier (Axon Instruments), acquired and analyzed with PatchMaster data acquisition software and using an Instrutech 1800 AD/DA board (HEKA Elektronik). Data acquisition was synchronized to the power supply controlling the micro-heater using an Arduino Uno. Pipettes for recording were pulled from borosilicate glass capillary and fire-polished to a resistance of 4-7 MΩ when filled with recording solution for inside- and outside-out recordings and 1-2 MΩ for whole-cell. Intracellular solutions in inside-out patches were changed using a custom built rapid solution changer. For whole-cell recordings all the bath solution was exchanged.

4 mM capsaicin stocks were prepared in ethanol, stored at −20°C and diluted to the desired concentration in recording solution as indicated before the experiments.

### Chemicals

Capsaicin and DTNB (5, 5′-Dithiobis (2-nitrobenzoic acid)) were purchased from Sigma-Aldrich. L-ANAP methyl ester (Methyl L-3-(6-acetylnaphthalen-2-ylamino)-2-aminopropionate) was purchased from AsisChem, Inc. (Waltham, MA).

### Temperature activation

Temperature activation was carried out using a micro-heater using the resistive heating principle as previously described (Islas et al., 2015), with the modification that, in this study, we used a programmable current-regulated DC power supply (Agilent) controlled via a Python program. For all recordings and temperature calibrations, the pipette with seal was placed approximately 27 µm from the micro-heater, in the case of whole-cell recording the cell was lifted and placed in front of the micro-heater. Before experiments, the temperature reached by the micro-heater was calibrated by the relation between resistance and temperature in an open pipette (Yao et al., 2009; Islas et al., 2015), which was obtained from an open pipette submerged in a chamber homogeneously heated by a peltier device and in which temperature was measured with a thermistor (Warner Instrument).

### Perfusion of cold solutions

For cooling experiments, the whole bath was exchanged by a bath solution cooled with ice and added sodium chloride, and gravity fed into the recording chamber. Temperature reached near the patch was measured with a thermistor.

### Cysteine modification

10 mM DTNB (5, 5′-Dithiobis (2-nitrobenzoic acid)) stocks were prepared in dimethyl sulfoxide, stored at -20°C, diluted to 10 µM in recording solution before experiments and used for just 2-3 hours.

### Incorporation of the fluorescent non-canonical aminoacid L-ANAP

L-ANAP incorporation into the construct rTRPV1-Y374TAG-mCherry was carried out using an aminoacyl tRNA synthetase and tRNA pair (RS-tRNA) contained in the pANAP vector, and optimized for mammalian cell expression (Chatterjee et al., 2013). This system incorporates the non-canonical aminoacid L-ANAP into an amber non-sense codon, TAG. HEK293 cells were transiently cotransfected with the pANAP vector containing the RS-tRNA pair and TRPV1-Y374TAG-mCherry in pcDNA3 with 6 µL of jetPei (PolyPlus). Total DNA concentration was 3 µg in 2 mL of culture medium. rTRPV1-Y374TAG-mCherryr and the RS-tRNA were prepared in a 3:1 proportion, respectively. L-ANAP was added 2 hours after plasmid transfection at a final concentration of 10 µM and cells were incubated for 48 hours for before deroofing and fluorescence analysis or whole-cell recording.

### Cell deroofing and fluorescence recording

Cell deroofing was carried out following a procedure slightly modified from (Heuser, 2000) and (Zagotta et al., 2016). HEK293 cells were plated in 35 mm glass bottom dishes (WPI) and transfected as indicated in the previous section. DMEM cell culture medium was removed and cells washed 3 times for 5 min, with 1.5 mL of stabilization buffer (KCl 70 mM, HEPES 30 mM, MgCl_2_ 1 mM, pH 7.4) plus CaCl_2_ 1mM, which is added just before it was used. Then, approximately 500 µL of poly-D-lysine solution (0.1 mM) resuspended in stabilization buffer, was used to completely cover the cells for 15 s. Next, cells were washed with 1.5 mL of hiposmotic solution (stabilization buffer diluted 1:2 with H_2_O) three times for 30 seconds, to remove poly-D-lysine and damaged cells. Finally 3.5 mL of stabilization buffer was added and a 0.5 s sonic pulse of approximately 15% of amplitude was applied using a Q-Sonica 50 W sonicator. If cells were not complete disrupted, a second or third pulse could be applied.

Membrane sheets left after deroofing were not easily visible in bright field and were identified by simultaneously measuring areas of fluorescence, containing emission from ANAP and mCherry. The custom fluorescence setup consists of a Nikon TE-2000u microscope coupled to an imaging spectrograph (Spectra pro 2150i, Acton instruments). Images were acquired using a CCD camera (iXon Ultra, Andor), with 300 msec exposure and gain set to 150. ANAP was excited with a 405 nm solid-state laser and mCherry was exited with the 514 nm line from an Argon-ion laser, using appropriate dichroic mirrors and filters. For ANAP quenching, a 20 µM dipicrylamide (DPA) solution was added, and quenching of ANAP fluorescence was followed by a multi-acquire image protocol in Micromanager software (Stuurman et al., 2007), acquiring an image every 5 seconds until stabilization of fluorescence decay (400-600 seconds). Membrane sheets thermal treatment was applied utilizing a specially designed aluminum chamber that receives a 35 mm glass-bottom petri dish in contact with a generic Multicomp 50 mm Peltier plate of 112.7 W. Temperature was calibrated and regulated by a CL-100 thermistor from Warner instruments. Temperature was changed immediately after cell deroofing and maintained at the desired value for 2 minutes, then the system was cooled down to room temperature and the ANAP quenching protocol was performed. Image analysis was done in Image J and IGOR software.

### Data analysis and mathematical modeling

All data were analyzed and plotted using Igor Pro v6 (Wavemetrics, Inc.). Pooled data are presented with the mean and standard deviation (s.e.m.). Statistical significance was assessed with a Student’s t-test as implemented in Igor pro. Significant differences between means were considered to exist went the p value was less than 0.01. Current activation by temperature was fit to a simple allosteric activation model to extract the enthalpy and entropy values (Scheme A in figure 4). The steady-state open probability as a function of temperature is given by the equation:

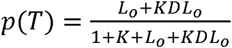

The temperature dependent equilibrium constant, K, given by:

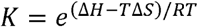

*L_o_* is the equilibrium constant for channel opening, and *D* is an allosteric coupling factor.

These two constants are set to be temperature-independent.

The predictions of the linear irreversible gating model (Figure 4d) were obtained by numerically solving the associated coupled equations with a Runge-Kutta fourth order method. The response to temperature ramps was simulated by simultaneously solving a linear equation for the temperature change. The numerical implementation was written in Igor Pro. The differential equations describing the model and the rate constants are as follow:

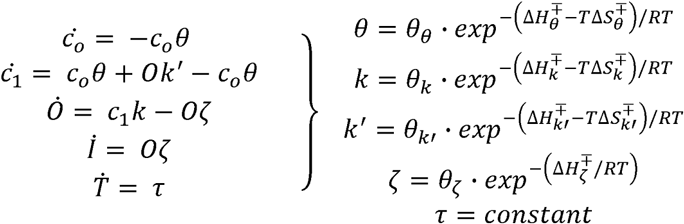

Here, the *θ_i_* are the diffusion-limited rate constants at 0K, 
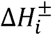
 and 
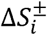
are the activation enthalpies and entropies, respectively, *R* is the gas constant and *T* is the temperature in Kelvin.

The temperature stimulus was chosen as a triangular symmetric ramp, which closely resembles de form of the stimulus provided by our microheaters. Initial conditions were T(0)=22°C, T(max)=chosen.

## Acknowledgments

We would like to thank Alejandra Llorente-Gil for expert technical help. This work was supported by grants from CONACYT CB-2015-252644 to L.D.I. and CB-2014-01-238399 to T.R., CONACYT-Fronteras de la Ciencia 77 to T.R. and L.D.I., DGAPA-PAPIIT-UNAM IN209515 to L.D.I. and IN200717 to T.R.

